# Real-world structure facilitates the rapid emergence of scene category information in visual brain signals

**DOI:** 10.1101/2020.03.24.004937

**Authors:** Daniel Kaiser, Greta Häberle, Radoslaw M. Cichy

## Abstract

In everyday life, our visual surroundings are not arranged randomly, but structured in predictable ways. Although previous studies have shown that the visual system is sensitive to such structural regularities, it remains unclear whether the presence of an intact structure in a scene also facilitates the cortical analysis of the scene’s categorical content. To address this question, we conducted an EEG experiment during which participants viewed natural scene images that were either “intact” (with their quadrants arranged in typical positions) or “jumbled” (with their quadrants arranged into atypical positions). We then used multivariate pattern analysis to decode the scenes’ category from the EEG signals (e.g., whether the participant had seen a church or a supermarket). The category of intact scenes could be decoded rapidly within the first 100ms of visual processing. Critically, within 200ms of processing category decoding was more pronounced for the intact scenes compared to the jumbled scenes, suggesting that the presence of real-world structure facilitates the extraction of scene category information. No such effect was found when the scenes were presented upside-down, indicating that the facilitation of neural category information is indeed linked to a scene’s adherence to typical real-world structure, rather than to differences in visual features between intact and jumbled scenes. Our results demonstrate that early stages of categorical analysis in the visual system exhibit tuning to the structure of the world that may facilitate the rapid extraction of behaviorally relevant information from rich natural environments.

## Introduction

In everyday situations, the input to our visual system is not random. It rather arises from highly organized scenes, which follow a predictable structure: In practically every real-word scene, visual information (such as the scene’s layout properties or the objects contained in a scene) is distributed in meaningful ways across space (Bar 2004; Kaiser et al., 2019a; Oliva & Torralba, 2007; Võ et al., 2019; Wolfe et al., 2011). Neuroimaging studies have shown that the visual system is sensitive to this structure, with cortical responses differing when scene elements do or do not adhere to typical real-world structure (Abassi & Papeo, 2019; Baldassano et al., 2017; Bilalic et al., 2019; Kaiser et al., 2014; Kaiser & Peelen, 2018; Kim & Biederman, 2011; Roberts & Humphreys, 2010). Although such studies suggest that the presence of real-world structure aids efficient scene representation, it is unclear how real-world structure impacts the representation of scene content: Specifically, does the presence of real-world structure facilitate the extraction of categorical information from a scene?

Evidence for an increase of visual category information in the presence of real-world regularities has already been reported for individual object processing. Several studies showed that typical real-world positioning enhances the neural representation of object category (Chan et al., 2010; de Haas et al., 2016; Kaiser & Cichy, 2018; Kaiser et al., 2018): for example, neural responses to an airplane are better discriminable from responses to other objects when the airplane is shown in the upper visual field, where it is typically encountered in the real world. Does the presence of real-world structure similarly facilitate the representation of categorical scene content in scenes?

To address this question, we used a jumbling paradigm (Biederman, 1972; Biederman et al., 1974) that manipulates natural scenes’ spatial structure: individual parts of the scene could either appear in their typical, “intact” positions or in atypical, “jumbled” positions (Figure 1). In a recent neuroimaging study (Kaiser et al., 2020a), we employed this paradigm to show that in scene-selective visual cortex (fMRI) and after 250ms of vision (EEG), spatially intact scenes were represented differently from jumbled scenes. Here, we analyzed the EEG data from this jumbling paradigm to investigate whether the typical real-world structure, in contrast to an atypical jumbled structure, facilitates the visual representation of scene category.

**Figure 1.**
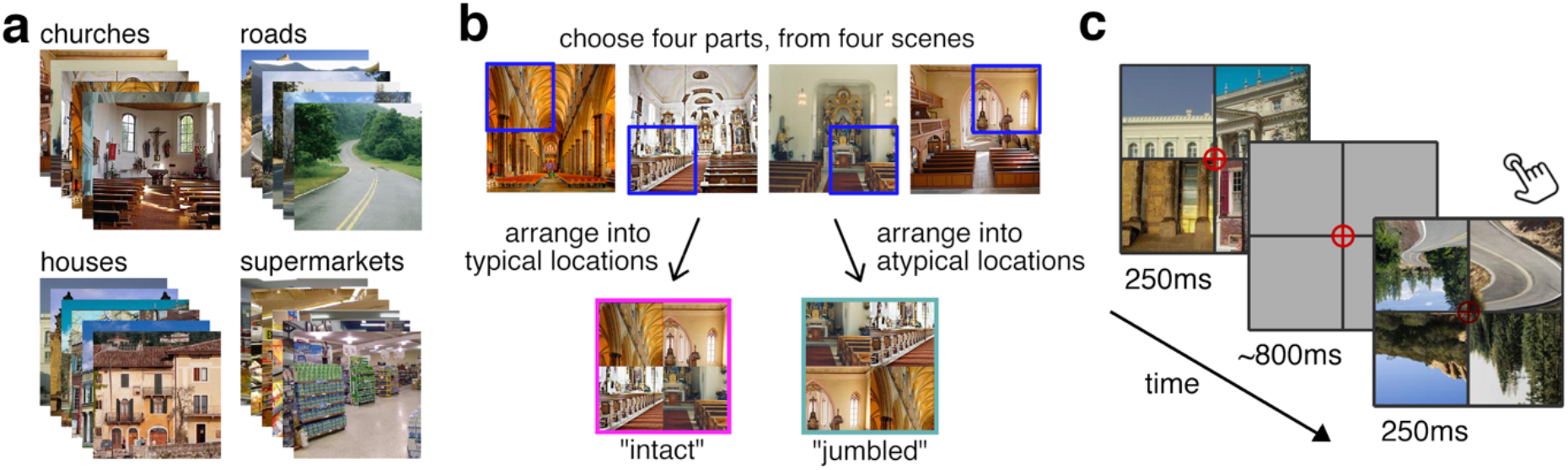
Experimental design. a) The stimulus set was constructed from natural scene photographs of four categories. b) Intact and jumbled scenes were created by combining parts of four different scenes of the same category in either typical locations or in locations (with positions swapped in a crisscrossed way). c) During the EEG experiment, participants viewed the scenes in upright and inverted orientation for 250ms each in random order. Participants performed an orthogonal task, where they responded whenever the fixation cross darkened.

To extract differences in category information between intact and jumbled scenes with high sensitivity, we used a cumulative multivariate decoding approach (Ramkumar et al., 2013), which maximizes the amount of data available at every time point along the processing cascade. In line with previous reports (Dima et al., 2018; Kaiser et al., 2019b, 2020b; Lowe et al., 2018), this analysis showed that scene category information emerges rapidly, within the first 100ms of vision. Critically, the early emergence of scene category information was facilitated for intact compared to jumbled scenes. This benefit was only present for upright, but not inverted scenes, indicating that the early facilitation of scene analysis is related to the presence of real-world structure, rather than differences in basic visual features.

## Materials and Methods

### Participants

Twenty healthy adults (mean age 26.6 years, *SD*=5.8; 9 female) participated. All participants had normal or corrected-to-normal vision. Participants provided informed consent and received either monetary reimbursement or course credits. All procedures were approved by the ethical committee of the Department of Psychology at Freie Universität Berlin and were in accordance with the Declaration of Helsinki.

### Stimuli

Stimuli were scenes from four different categories: churches, houses, roads, and supermarkets (Figure 1a). The stimuli were taken from an online resource (Konkle, Brady, Alvarez, & Oliva, 2010). For each category six different exemplars were used. To manipulate scenes’ adherence to real-world structure, we first split each original image into quadrants. We then systematically recombined quadrants from different scenes such that the scenes’ spatial structure was either intact or jumbled (Figure 1b). For the intact scenes, four fragments from four different scenes of the same scene category were combined in their correct spatial locations. For the jumbled scenes, four fragments from four different scenes of the same scene category were combined, but their spatial locations were arranged in a crisscrossed way. This jumbling manipulation simultaneously disrupted multiple structural regularities in the scene, such as visual feature distributions, scene geometry, absolute and relative object positions, and cues to 3D-structure. Additionally, the stimulus set entailed scenes that were jumbled in their categorical content (with the individual scene parts stemming from different categories); these scenes were created to answer a different research question (see Kaiser et al., 2020a) and not used in the analyses reported in this paper. In both conditions relevant for this paper, we used fragments from four different scenes to equate the presence of visual discontinuities between fragments. Separately for each participant, 24 unique intact and 24 unique jumbled stimuli were generated by randomly drawing suitable fragments from different scenes. Each scene was presented upright and upside-down.

### Paradigm

During the EEG experiment, the different stimuli were randomly intermixed within a single session. Within each trial, a scene appeared for 250ms. Stimuli appeared in a black grid (4.5deg visual angle), which served to mask visual discontinuities between quadrants (Figure 1c). Each trial was followed by an inter-trial interval which randomly varied between 700ms and 900ms. For this paper, only parts of the collected data – spatially intact and spatially jumbled scenes in upright and upside-down orientation – were analyzed. Each of these four conditions covered 384 trials (equating to 96 trials per scene category). Additionally, 1,152 target trials were measured. During the target trials the crosshair changed into a slightly darker red at the same time the scene was presented. When detecting a target, participants had to press a button; additionally, they were asked to blink during the target trials, making it easier for them to refrain from blinking during non-target trials. Target detection was purposefully made challenging to ensure sufficient attentional engagement (mean accuracy 78.1%, *SE*=3.6%). Target trials were not included in the subsequent analyses. Furthermore, 1,536 trials where the scene’s categorical structure was altered have been recorded. This data has been analyzed elsewhere (see Kaiser et al., 2020a). Further, participants were instructed to maintain central fixation throughout the experiment. Stimulus presentation was controlled using the Psychtoolbox (Brainard, 1997).

### EEG recording and preprocessing

The EEG data were the same as in Kaiser et al. (2020a). EEG signals were recorded using an EASYCAP 64-electrode system and a Brainvision actiCHamp amplifier. For two participants, due to technical problems, only data from 32 electrodes was recorded. Electrodes were arranged in accordance with the 10-10 system. EEG data was recorded at 1000Hz sampling rate and filtered online between 0.03Hz and 100Hz. All electrodes were referenced online to the Fz electrode. Offline preprocessing was performed using FieldTrip (Oostenveld et al., 2011). EEG data were epoched from −200ms to 800ms relative to stimulus onset, and baseline-corrected by subtracting the mean pre-stimulus signal. Channels and trials containing excessive noise were removed based on visual inspection. Blinks and eye movement artifacts were removed using independent components analysis and visual inspection of the resulting components (Jung et al. 2000). The epoched data were down-sampled to 200Hz.

### EEG decoding

Decoding analyses were performed using CoSMoMVPA (Oosterhof et al., 2016). To track cortical representations across time, we used a cumulative classification approach that for each time point across the epoch takes into account all time points prior to the current time point (Ramkumar et al., 2013). This classification technique uses larger amounts of data at each subsequent time point, while maintaining temporal precision in the forward direction (i.e., it only collapses across information backwards in time, but not forwards). Cumulative decoding may thus provide increased sensitivity for detecting decoding onsets, compared to standard timeseries decoding (Grootswagers et al., 2017).

We used such cumulative classifiers to discriminate between the four scene categories. This analysis was done separately for the intact and jumbled scenes. Classification analyses were performed repeatedly, with the amount of information available to the classifier accumulating across time (Figure 2). That is, for the first time point in the epoch, the classifier was trained and tested on response patterns across the electrodes at this time point. At the second time point in the epoch, the classifier was trained and tested on response patterns across the electrodes at the first and second time point in this epoch. Finally, at the last time point in the epoch, the classifier was trained on response patterns across all electrodes and at all time points in this epoch.

**Figure 2.**
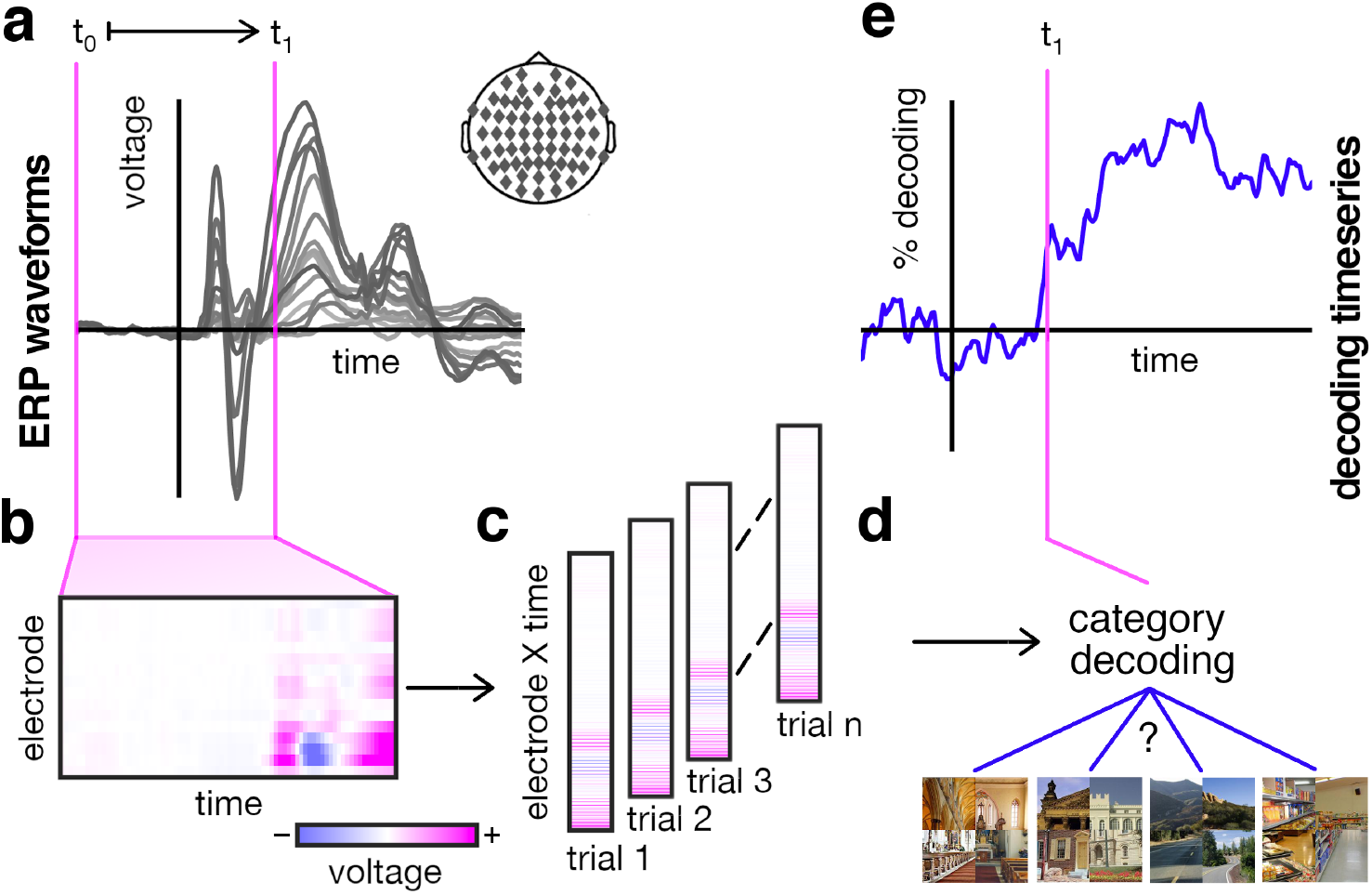
Schematic depiction of the cumulative decoding approach. a) For each time point t_1_ across the epoch, a separate decoding analysis was performed. b) For each of these analyses, we aggregated ERP waveforms across all EEG electrodes and all time points between t_1_ and the beginning of the epoch (t_0_). c) For each trial, we then unfolded these two-dimensional response pattern across electrodes and time into a one-dimensional response pattern. d) These one-dimensional response patterns were first subjected to PCA analysis to reduce dimensionality (see Materials and Methods) and then fed to LDA classifiers, which were trained to discriminate the four scene categories. Decoding accuracy was computed by repeatedly assessing classifier performance on single trials left out during classifier training. e) Repeating this analysis across time yielded a decoding timeseries with 200Hz resolution. Importantly, the cumulative nature of this analysis allowed us to increase power by increasing the amount of data available to the classifier without losing temporal precision regarding the onset of category information.

The richer information contained in these cumulative response patterns comes at the expense of a higher dimensionality of the data, which potentially harms classification. To reduce the dimensionality of the data at each time point, we performed principal component analyses (PCAs). These PCAs were always done on the classifier training set and the PCA solution was projected onto the testing set (Grootswagers et al., 2017). For each PCA, we retained as many components as needed to explain 99% of the variance in the training set data (average number of components retained at example time points; at 0ms: 225, SE=11; at 200ms: 250, SE=10; at 800ms: 269, SE=10).

For classification, we used linear discriminant analysis (LDA) classifiers. For each classifier, the covariance matrix was regularized by adding the identity matrix scaled by one percent of the mean of the diagonal elements (as implemented in the *cosmo_classify_lda* function in CoSMoMVPA; Oosterhof et al., 2016). Classification was performed in a cross-validation scheme with 12 distinct folds. Classifiers were trained on data from 11 of these folds and tested on data from the left-out fold. The amount of data in the training set was always balanced across the four categories. Classification was done repeatedly until every fold was left out once. Classification accuracies were averaged across these repetitions. These analyses resulted in separate decoding timeseries for intact and jumbled scenes, which reflect the temporal accrual of category information (i.e., how well the four categories are discriminable from the neural data).

### Statistical testing

To compare decoding timeseries against chance level and the different conditions’ decoding timeseries against each other, we used a threshold-free cluster enhancement (TFCE) procedure (Smith & Nichols, 2009). Multiple-comparison correction was based on a sign-permutation test (with null distributions created from 10,000 bootstrapping iterations) as implemented in CoSMoMVPA (Oosterhof et al., 2016). The resulting statistical maps were thresholded at *z*>1.96 (i.e., *pcorr<.05*). However, the onset of statistical significance for TFCE methods may be biased by the presence of strong clusters following the onset (as expected from the cumulative decoding performed here) and can therefore not be directly interpreted (Sassenhagen & Draschkow, 2019). We thus additionally provide statistics for conventional one-sample t-tests, which we corrected for multiple comparisons using false-discovery-rate (FDR) corrections. For all tests, only clusters of at least 4 consecutive significant time points (i.e., more than 20ms) were considered.

### Data availability

Data are publicly available on OSF (doi.org/10.17605/OSF.IO/ECMA4).

## Results

We first analyzed data from the upright scenes, where we expected a facilitation of category information for spatially intact, compared to jumbled, scenes. We found that EEG signals conveyed robust scene category information: categories were discriminable for both intact scenes (significant decoding obtained by TFCE statistics: between 75ms and 800ms; significant decoding obtained by FDR-corrected statistics: between 75ms and 800ms) and jumbled scenes (TFCE: between 120ms and 800ms; FDR: between 135ms and 800ms) (Figure 3a). Crucially, we found significantly enhanced decoding for the spatially intact scenes, compared to the jumbled scenes (TFCE: between 105ms and 800ms; FDR: between 105ms and 800ms) (Figure 3c).

**Figure 3.**
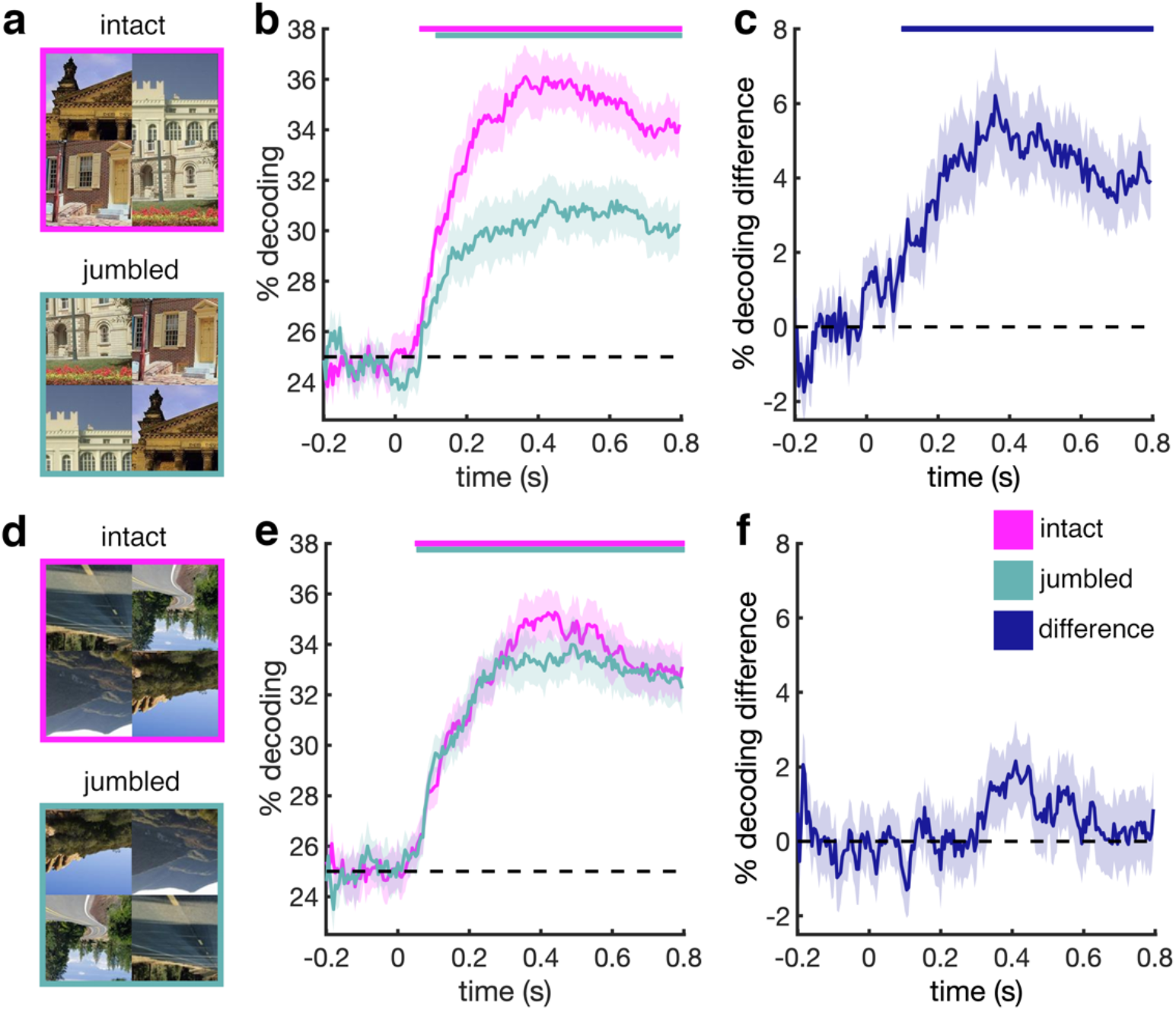
Decoding of scene category for intact and jumbled scenes. a) First, we decoded the category of intact and jumbled scenes when they were presented upright. b) This analysis revealed widespread clusters of category decoding for both intact and jumbled scenes. c) Critically, we found more accurate decoding of scene category when the scene was intact, suggesting that adherence to real-world structure boosts early visual category information. d) Second, we decoded the category of upside-down scenes. e) For upside-down scenes, category could be similarly decoded from the EEG signals. f) However, there was no benefit of intact scene structure when the scenes were inverted, suggesting that adherence to real-world structure, rather than low-level differences, explain the enhanced category decoding for structured scenes when they are upright. Error margins indicate standard errors of the difference. Significance markers indicate p<0.05, corrected for multiple comparisons using TFCE.

The inclusion of inverted scenes allowed us to investigate whether the effects of scene structure were genuinely related to the scenes adhering to real-world structure, rather than differences in their low-level visual attributes. If the enhanced category information for spatially intact scenes is indeed related to their adherence with real-world structure, then no effects should be seen when the same scenes are viewed upside-down, as all inverted scenes do not adhere to real-world structure.

Performing the category decoding analysis on the inverted scenes (Figure 3d) revealed a qualitative difference to the upright scenes: the effect of scene structure was significantly stronger for the upright scenes (TFCE: between 170ms and 800ms; FDR: between 95ms and 115ms, and between 185ms and 800ms). Indeed, no significant differences between intact and jumbled scenes were observed for the inverted scenes, although the category of both intact scenes (TFCE: between 55ms and 800ms; FDR: between 60ms and 800ms) and jumbled scenes (TFCE: between 60ms and 800ms; FDR: between 75ms and 800ms) could be decoded from the EEG signals (Figure 3e/f). This indicates that the early facilitation of scene category information for spatially structured scenes can be attributed to the scenes adhering to typical real-world structure, rather than to low-level features differing between the intact and jumbled scenes.

Our results establish that for processing of upright scenes, scene structure matters more than for processing inverted scenes. Additionally, one can also ask how robustly category information emerges as a function of whether the scene is presented upright or upside down. To answer this question, we directly compared category information for the intact upright scenes, the jumbled upright scenes, and the inverted scenes (Figure 4a); as for the inverted scenes we found no statistical differences between the intact and jumbled scenes, we averaged across them. We found that category decoding accuracy for the inverted scenes was numerically in between the intact and jumbled upright scenes (Figure 4b). When directly comparing the decoding time courses (Figure 4c), we found that category decoding was not significantly stronger in the intact upright scenes, compared to the inverted scenes. By contrast, category decoding for the jumbled scenes was significantly weaker than for the inverted scenes (TFCE: between 170ms and 800ms; FDR: between 200ms and 800ms). This result suggests that for the inverted scenes, category can be decoded similarly as for the intact upright scenes. However, once the structure of an upright scene is destroyed, only weaker categorical representations emerge in the visual system.

**Figure 4.**
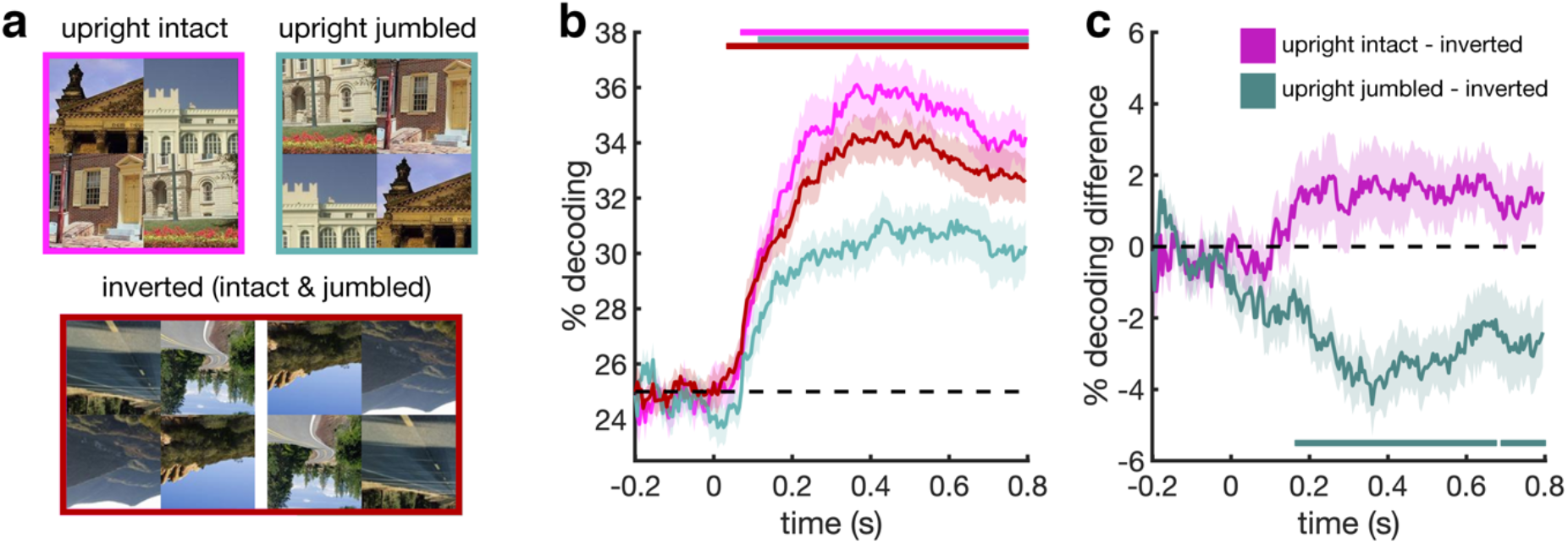
Comparing category decoding between upright and inverted scenes. a) We compared the emergence of category information for the intact upright scenes, the jumbled upright scenes, and the inverted scenes; for the inverted scenes, we averaged across the intact and jumbled scenes, as no significant differences between the two were found. b) Numerically, category decoding accuracy for the inverted scenes was in between the accuracies observed for the intact and jumbled upright scenes. c) When subtracting decoding in the inverted condition from decoding in the upright conditions, we found that statistically, category information was comparable for intact upright scenes and inverted scenes. By contrast, weaker category information was found for the jumbled upright scenes, compared to the inverted scenes, suggesting that jumbling specifically harms the emergence of category information in upright scenes. Error margins indicate standard errors of the difference. Significance markers indicate p<0.05, corrected for multiple comparisons using TFCE.

## Discussion

Our results provide evidence that real-world regularities facilitate the extraction of scene category information during visual analysis. We show that this facilitation of category information emerges within the first 200ms of vision. Our findings highlight the pervasive role of real-world structure in perceptual processing, suggesting that already at relatively early processing stages cortical scene representations are tightly linked to the typical composition of our daily surroundings.

Here, we used a cumulative decoding technique to establish differences in the initial emergence of information in EEG signals. This technique uses all the available historical data (i.e., data prior to the current time point) for classification. Together with using PCA for dimensionality reduction, the availability of this larger amount of data promises high detection sensitivity. The availability of historical data at later time points may also hold true for the brain, where downstream regions have access to information coded earlier in upstream regions. However, as a note of caution, classifiers may also use temporally distinct information that is not necessarily available in the same way in the brain, particularly when looking at late processing stages. Cumulative decoding nonetheless provides a useful approach to reveal early differences in cortical information processing.

The early facilitation of category information is consistent with results from single-object processing, where representations of individual objects are rapidly enhanced – within the first 150ms of vision – when the objects appear in their typical real-world locations, such as an eye in the upper visual field (Issa & DiCarlo, 2012) or a shoe in the lower visual field (Kaiser et al., 2018). Together, these findings therefore support the idea that real-world structure can boost basic visual analysis across diverse stimuli and processing levels (Kaiser et al., 2019a).

When directly comparing neural category information in upright and inverted scenes, we found that it was equally pronounced when the scenes were intact and upright and when the scenes were inverted, regardless of their structural arrangement – only when the upright scenes were jumbled, we found significantly reduced category information. One interpretation of this result is that jumbling causes a specific disruption for upright scenes, as for these scenes the jumbling manipulating may be perceptually more salient. Alternatively, the pattern of results may be explained by an interaction of two different effects: The inverted intact scenes still retain the intact relative positioning of their parts, which may explain why they are better decodable than the upright jumbled scenes. The inverted jumbled scenes do not have this intact relative positioning, but by means of inversion they gain an intact absolute positioning of their parts (e.g., a piece of sky would be in the upper part of an inverted jumbled scene, which is where it belongs); this may explain why these scenes yield better category decoding than upright inverted scenes. At this point, further research is needed to fully understand this pattern of results.

While our effects demonstrate an enhanced early representation of scenes that adhere to real-world structure compared to scenes that do not, studies on object-scene consistency suggest that EEG waveforms only become affected by typical object positioning after around 250ms of vision (Coco et al., 2019; Draschkow et al., 2018; Ganis & Kutas, 2003; Mudrik et al., 2010, 2014; Võ & Wolfe, 2013). How do these early and late effect of scene structure relate to each other?

As one possibility, later effects may partly reflect increased responses to inconsistencies, rather than an enhanced processing of consistent scene-object combinations (Faivre et al., 2019). Together with our results, these findings may suggest that early responses are biased towards scenes that predictably follow real-world structure, whereas later responses may be more biased towards violations of this structure. This idea is consistent with a recent proposal in predictive processing, which suggests a temporal succession of more general processing biases, first towards the expected and then towards the surprising (Press et al., 2020).

Alternatively, the beneficial effects of real-world regularities may not immediately result in consistency signals: Whether visual inputs generally are consistent with our real-world experience may only be analyzed following more basic visual analysis. Supporting this idea, generic consistency signals in our data only emerge later than the enhanced category processing: As previously reported, intact and jumbled scenes (independent of their category) evoked reliably different responses only after 255ms of processing (Kaiser et al., 2020a).

More broadly, the findings can add to our understanding of efficient everyday vision. Even under challenging real-world conditions, human vision is remarkably efficient – in fact, much more efficient than findings from simplified laboratory experiments would predict (Wolfe et al., 2011; Peelen & Kastner, 2014). Behavioral studies using jumbling paradigms have suggested that typical scene structure contributes to this efficiency: when scenes are structurally intact, observers can better categorize them (Biederman et al., 1974), recognize objects within them (Biederman, 1972), or detect visual changes in the scene (Varakin & Levin, 2008). These perceptual benefits may be linked the rapid facilitation of neural category information for typical scenes observed in the current study. However, our participants performed an orthogonal fixation task, which precludes directly linking brain and behavior here. Future studies combining neural recordings with naturalistic behavioral tasks may reveal that the early cortical tuning to real-world structure may be a crucial asset for solving complex real-world tasks.

## Acknowledgements

We thank Sina Schwarze for help in EEG data collection. D.K. and R.M.C. are supported by Deutsche Forschungsgemeinschaft (DFG) grants (KA4683/2-1, CI241/1-1, CI241/3-1). R.M.C. is supported by a European Research Council Starting Grant (ERC-2018-StG 803370). G.H. is supported by a PhD fellowship of the Einstein Center for Neurosciences Berlin.

## Notes

### Competing Interest Statement

The authors have declared no competing interest.

